# Serum Flt3 ligand is biomarker of progenitor cell mass and prognosis in acute myeloid leukemia

**DOI:** 10.1101/588319

**Authors:** Paul Milne, Charlotte Wilhelm-Benartzi, Michael R. Grunwald, Venetia Bigley, Amy Publicover, Sarah Pagan, Helen Marr, Gail Jones, Anne Dickinson, Angela Grech, Alan Burnett, Nigel Russell, Mark Levis, Steven Knapper, Matthew Collin

## Abstract

Fms-like tyrosine kinase 3 (Flt3) is a hematopoietic growth factor receptor expressed on lymphomyeloid progenitors and frequently, by AML blasts. Its ligand, Flt3L, has non-hematopoietic and lymphoid origins, is detectable during homeostasis and increases to high levels in states of hypoplasia due to genetic defects or treatment with cytoreductive agents. Measurement of Flt3L by ELISA reveals that Flt3^+^AML, is associated with depletion of Flt3L to undetectable levels. After induction chemotherapy, Flt3L is restored in patients entering CR, but remains depressed in those with refractory disease. Weekly sampling reveals marked differences in the kinetics of Flt3L response during the first 6 weeks of treatment, proportionate to the clearance of blasts and cellularity of the BM. In the UK NCRI AML17 trial, Flt3L was measured at day 26 in a subgroup of 135 patients with Flt3 mutation randomized to the tyrosine kinase inhibitor lestaurtinib. In these patients, attainment of CR was associated with higher Flt3L at day 26 (Mann-Whitney p < 0.0001). Day 26 Flt3L was also associated with survival: Flt3L ≤ 291pg/ml was associated with inferior event-free survival; and, Flt3L >1185pg/ml was associated with higher overall survival (p = 0.0119). Serial measurement of Flt3L in patients who had received a hematopoietic stem cell transplant for AML further illustrated the potential value of declining Flt3L to identify relapse. Together these observations suggest that measurement of Flt3L provides a non-invasive estimate of progenitor cell mass in most patients with AML, with the potential to inform clinical decisions.

**Graphical abstract:** 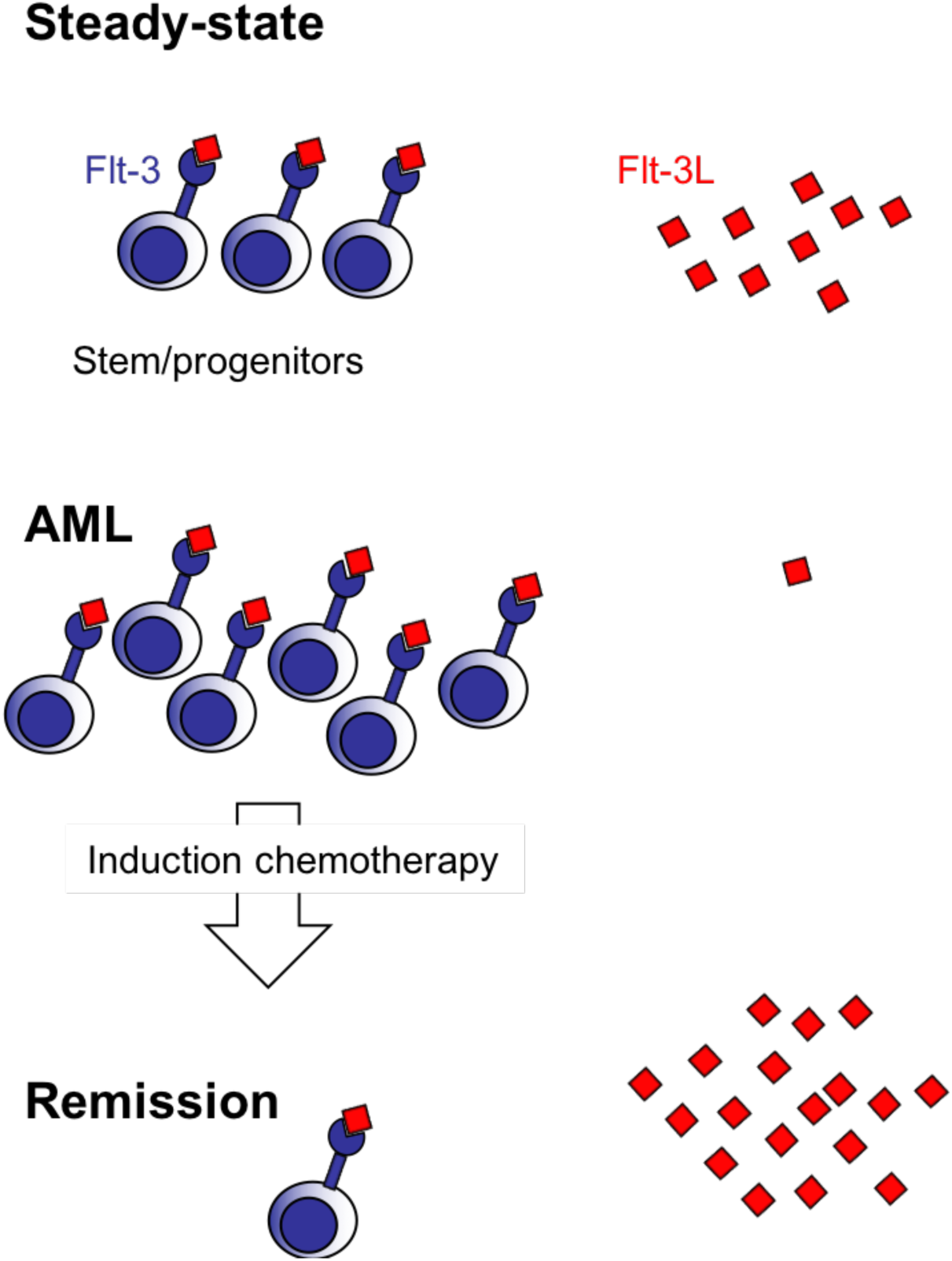

## Introduction

Acute myeloid leukemia is a high-risk malignancy with fewer than one-third of patients surviving at 5 years^1, 2^. Despite intensive chemotherapy, therapeutic resistance and relapse are the main causes of treatment failure. The early identification of primary treatment refractoriness is especially challenging and independent of standard clinical, cytogenetic and molecular risk factors ^3^, even with integrated molecular genetic information ^4^. Large multi-cohort studies show that up to 30% of younger patients have primary refractory disease or relapse-free survival (RFS) ≤3 months, rising to 40% with RFS ≤6 months and 57% with RFS ≤ 1 year ^3^. A limited ability to predict a poor therapeutic response, in spite of advances in genetic risk profiling and minimal residual disease detection, continues to impede the delivery of risk-adapted therapy ^5–7^. Salvage after relapse is also disappointing with fewer than 50% achieving a subsequent complete remission (CR), leading to median survivals of less than 1 year ^8^. Outcomes are considerably worse for patients older than 60, who constitute the majority of cases ^9^.

Fms-like tyrosine kinase-3 ligand (Flt3L) is a growth factor involved in the maintenance and proliferation of hematopoietic stem and progenitor cells ^10^. Its receptor Flt3 (CD135), is a tyrosine kinase expressed predominantly on primitive CD34^+^CD38^low^progenitor fractions ^11^. Several lines of evidence indicate that Flt3L rises when the hematopoietic stem cell compartment is compromised. Flt3L is increased in Fanconi anemia, acquired aplastic anemia, chemotherapy or radiotherapy-induced aplasia, and heterozygous GATA2 mutation ^12–15 16 17^. Elevated Flt3L was also observed in patients with AML at day 15 of treatment and shown to be a potential mechanism of resistance to therapy with Flt3 inhibitors ^10^.

Conversely, suppression of Flt3L in association with expansion of hematopoietic progenitors, has not been reported. Previous studies investigating hypercellularity associated with myelodysplastic syndrome (MDS) lacked a sufficiently sensitive ELISA to determine levels much below the normal range ^18^. Mobilization of peripheral blood stem cells (PBSC) with G-CSF is paradoxically associated with a slight elevation of Flt3L ^19, 20^. An inverse relationship between Flt3L and cells bearing its receptor might be predicted from the molecular control of other elements of hematopoiesis such as platelets ^21^.

Here we show that the presence of CD135+ AML blasts is associated with significant depression of Flt3L and that measurement of this biomarker may have utility in diagnosis, early assessment of response to therapy and monitoring of remission status.

## Methods

### Single centre patients and controls

Patients with heterozygous mutation of *GATA2*, aplastic anemia, MDS and AML were recruited from the Northern Centre for Cancer Care, Freeman Hospital, Newcastle upon Tyne Hospitals NHS Foundation Trust under ethical permission from the Newcastle and North Tyneside Research Ethics Committee 1 Reference 08/H0906/72 (Cohort A). Informed consent was obtained in accordance with the Declaration of Helsinki. Additional patients with AML were recruited from Johns Hopkins Sydney Kimmel Comprehensive Cancer Center, Baltimore under the IRB-approved protocol J10145. A total of 84 patients were recruited at Newcastle, UK and 8 at Johns Hopkins, Baltimore (Cohort B). All but 2 patients received intensive induction chemotherapy with cytarabine in combination with either daunorubicin, etoposide or mitoxantrone. Details of all patients included in the study under these protocols are summarized in Table S1. Healthy control blood was obtained from volunteer donors.

### AML17 patients

The UK NCRI AML 17 study (ISRCTN 55675535) was a large, prospective phase 3 multicenter trial for patients with newly diagnosed AML or high-risk MDS (≥10% marrow blasts) that ran 2009-2014 at more than 130 centers in the United Kingdom, Denmark, and New Zealand, recruiting 1206 patients ^22, 23^. 325 patients with Flt3 mutation were randomized 2:1 to receive lestaurtinib (CEP701); 140 of those randomized provided a blood sample at day 26 for measurement of Flt3 ligand.

### Flt-3L ELISA

Serum Flt-3L was measured with quantikine human Flt3/Flk-2 ligand immunoassay, according the manufacturer’s instructions (R&D Systems).

### Flow cytometry

Bone marrow or peripheral blood blasts were prepared with ficoll gradients and stained in aliquots of approximately 1 million cells. Antibodies were from BD Biosciences (www.bdbiosciences.com) unless stated otherwise and are denoted as: antigen fluorochrome (clone). CD3 FITC (SK7); CD10 PE-TR (B-Ly6; Beckman Coulter; www.beckmancoulter.com); CD14 FITC (M5E2); CD16 FITC (NKP15/Leu-11a); CD19 FITC (4G7); CD20 FITC (L27); CD34 APCCy7 (581); CD38 PECy7 (HB7); CD45 V450 (2D1); CD45RA BV510 (HI100; Biolegend); CD56 FITC (NCAM16.2); CD90 PerCPCY5.5 (5E10; Biolegend); CD123 PerCPCY5.5 (7G3); CD135 PE (4G8); HLA-DR V500 (L243/G46-6). Events were acquired with a B-D Biosciences Fortessa X20 running Diva version 8 and analyzed with Treestar FlowJo version 7.6.5. Blasts were identified as a high FSC/SSC lineage-negative population and Median Fluorescence Intensity (MFI) of CD135 was calculated by subtraction of an appropriate isotype control.

### Statistical Analyses

Figures were plotted in R (version 3.0.1), http://www.r-project.org/. or Prism version 7.0d (GraphPad Software Inc). All statistical analyses for AML17 patient data were carried out in R 3.0.1. Date of death was used to calculate overall survival (OS). Event-free survival (EFS) events were defined as: failure to achieve CR, disease relapse or death from any cause. For both survival endpoints, patients without a date of relapse or death were censored at the date of last visit or the date at which patients were last known to be alive. Remission status was measured using seven categories namely: Complete remission (CR), Death before chemotherapy started, Induction death (ID), Resistant disease (RD), Partial remission (PR), CR with incomplete count recovery (CRi), and Other. Remission status was then dichotomized into CR versus non-CR. The optimal cut point separating patients in CR from those with no CR was defined by the Youden method using the Receiver Operating Characteristic (ROC) analysis in the R package ‘Optimal Cutpoints’ ^24^. We then plotted a ROC curve for this cut point and determined the area under the curve (AUC), sensitivity and specificity. The R package “Survminer” was then used to determine the optimal cut point for overall and event-free survivals based on the maximally selected log-rank statistic. The two survival curves split by this cut point were then plotted using the descriptive Kaplan Meier method with the log-rank p-value.

## Results

### Association of Flt3L with progenitor cell mass and Flt3 expression in AML

The relationship between Flt3L level and progenitor cell mass was explored in a number of distinct patient groups recruited at the Newcastle center, including Cohort A patients with AML (**Figure 1A**). As reported previously, patients with hypoplasia of the hematopoietic stem/progenitor compartment due to aplastic anemia or germline *GATA2* mutation, have very elevated serum Flt3L ^12, 13, 16, 17^. A cohort of ambulatory patients with MDS showed increased variance of Flt3L levels while patients admitted for treatment of AML mostly had low or undetectable levels of Flt3L. A potential link between decreasing Flt3L and progression to AML was suggested by the observation that the two MDS patients with the lowest Flt3L (≤7pg/ml) progressed to AML within 6 months. Of 75 consecutive patients admitted for treatment of AML, 71 had Flt3L below the level of healthy controls (range 48.3 - 173.8 pg/ml; n = 20) and 63 below the limit of detection of 7pg/ml. Further clinical details of patients with AML are given in **Table S1.**

**Figure 1.**
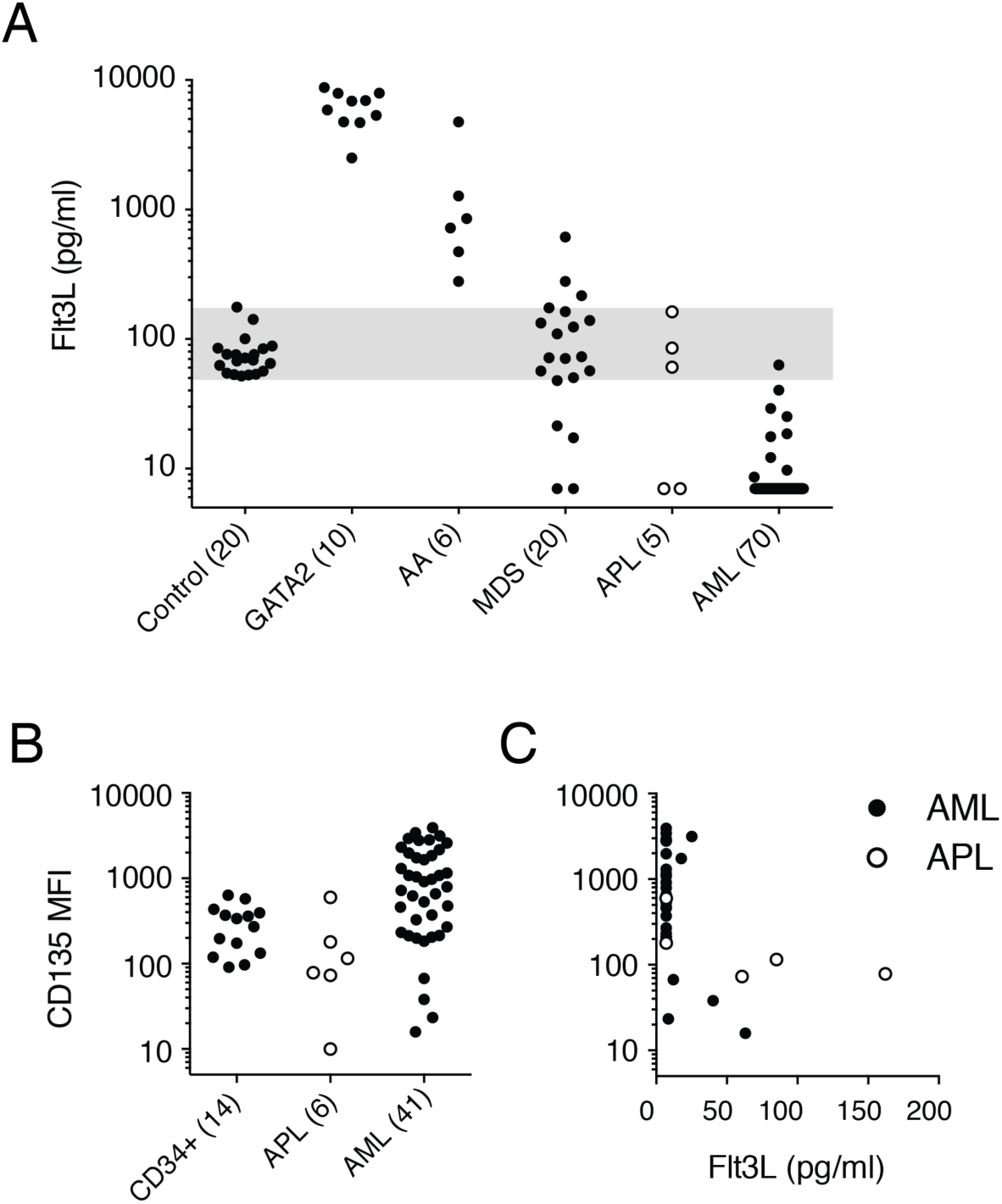
Association of Flt3L with progenitor cell mass and Flt3 expression in AML Cohort A. **A**.Serum Flt3L measured by ELISA (limit of detection 7pg/ml) in healthy controls (range 48.3 - 173.8 pg/ml), individuals with heterozygous germline GATA2 mutation, acquired aplastic anemia (AA), myelodysplastic syndrome (MDS), acute promyelocytic leukemia (APL; open symbols) and acute myeloid leukemia (AML). Numbers in each cohort indicated in parentheses. **B**.Level of Flt3 (CD135) expression indicated by median fluorescence intensity (MFI; minus isotype control) in control CD34+ BM progenitors, APL blasts and AML blasts. Numbers in each cohort indicated in parentheses. Note MFI is a log scale. **C**.Inverse relationship between CD135 expression and Flt3L in 14 AML and 4 APL cases in which both measurements were performed.

Clinical variables were explored in order to try to understand the reason for Flt3L suppression in most but not all cases of AML. There were no statistically significant relationships between WBC at diagnosis, % BM blasts, cytogenetics, presence of Flt3 ITD, blast phenotype and disease stage (**Figure S1**). The only notable feature was an association between the diagnosis of acute promyelocytic leukemia (APL) or the presence of t(15;17) in patients with normal Flt3L. There were 5 cases of APL of which 3 had normal Flt3L, compared with only 1 of 70 cases of non-APL.

The expression of Flt3 (CD135) was analysed by flow cytometry in a subset of patients shown in A, including the 5 patients with APL. This confirmed that Flt3 was expressed at a lower level on APL than most AML blasts (**Figure 1B**) and that there was an approximate inverse relationship between the expression of Flt3 and the level of Flt3L across all cases (**Figure 1C**).

### Association of Flt3L with remission assessment post-chemotherapy

Further samples, following two rounds of intensive chemotherapy, were available from 28 patients in Cohort A (Newcastle) who had undetectable Flt3L at diagnosis. These were taken at the time of remission assessment, between 28-42 days after the start of each round of chemotherapy. Fifteen patients were in morphological CR after both courses of chemotherapy and showed recovery of Flt3L to normal or supra-normal levels at both time points (**Figure 2A**). A further 8 patients did not achieve morphological CR after the first course but subsequently entered remission after the second course (D+A or Fla-Ida salvage) (**Figure 2B**). In contrast, Flt3L remained suppressed in 5 patients who did not enter CR after 2 cycles of chemotherapy. In addition, 3 patients thought to be in morphological CR after the first course failed to show any appreciable recovery of Flt3L after either course 1 or course 2. These patients all relapsed within 6 months of diagnosis at 114-172 days, compared with RFS of 317-825 days for patients in A and B, whose Flt3L was elevated after treatment (**Figure 2C**).

**Figure 2.**
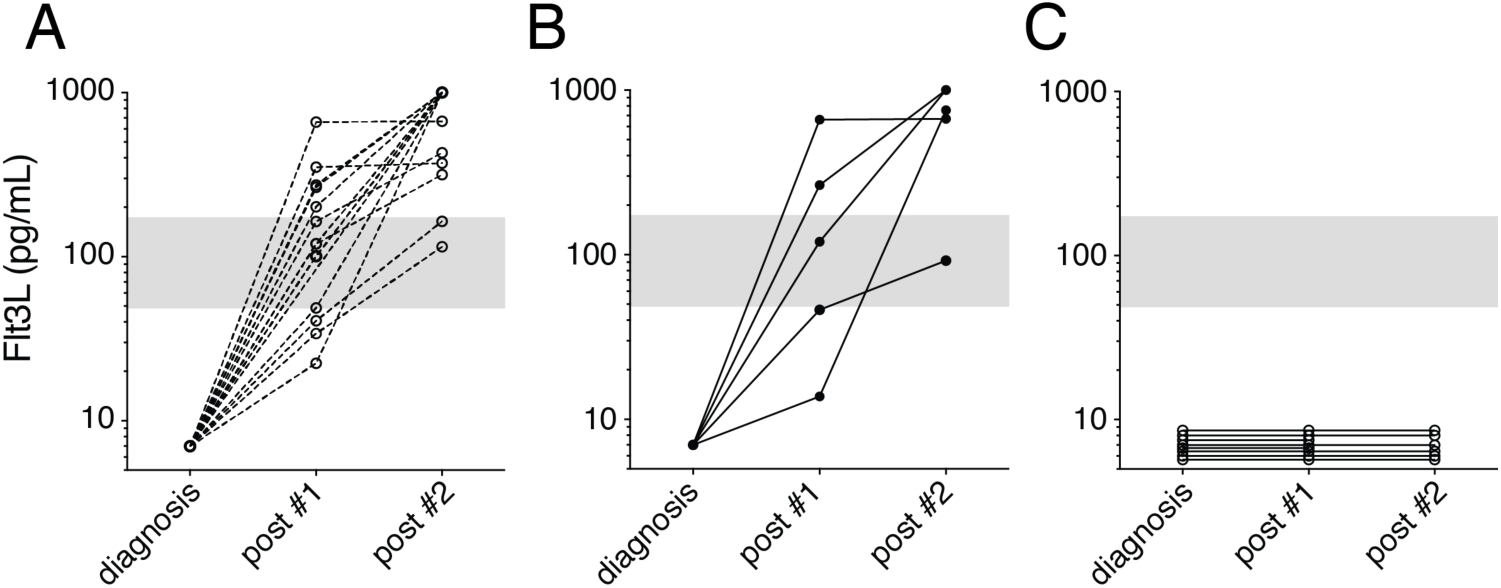
Serial serum Flt3L measurements in AML patients. **A** Patients were sampled at diagnosis and following each of 2 cycles of intensive chemotherapy with D+A (3+10) or FLA-Ida. (Cohort A, Newcastle). Patients who entered continuing morphological CR after 2 courses maintained increasing Flt3L at remission assessments performed 28-42 days after the start of chemotherapy (range of detection 7-1000pg/ml; n = 15; broken lines). **B**.Patients who did not achieve morphological CR after the first course of chemotherapy but subsequently entered remission after the second, also demonstrated increasing Flt3L over both cycles (n = 5; filled circles and solid lines) **C**.Patients who remained refractory after 2 courses of chemotherapy had undetectable Flt3L (n = 5; open circles and solid lines). The graph also includes a further 3 patients thought to be in morphological CR after the first course of chemotherapy who experienced relapse within 6 months.

### Kinetics of Flt3L response during induction chemotherapy

The preceding data suggest that the level of Flt3L was highly discriminatory between a state of normal hematopoiesis and the presence of CD135^+^AML blasts. Undetectable Flt3L appeared to associate with resistant or refractory disease when measured between courses of chemotherapy at the time of remission assessments by bone marrow aspiration. Weekly sampling was then performed in a subset of 12 non-APL AML patients (4 Cohort A and 8 Cohort B) in order to determine whether it might be possible to predict remission status from the kinetics of Flt3L response. Six patients who eventually achieved CR showed Flt3L elevation of 1,000-4,000 pg/ml, peaking between day 14 and 21 and returning to the normal range after chemotherapy (**Figure 3A**). In contrast, 4 patients who did not enter CR showed more modest elevations of less than 1,000 pg/ml that subsequently dropped back to below normal or undetectable levels at day 42 (**Figure 3B)**. Two patients in the series remained pancytopenic after chemotherapy induction and were found to be have hypocellular bone marrows without detectable blasts. In these cases, Flt3L remained high (>1000pg/ml) (**Figure 3C).** One was salvaged by hematopoietic stem cell transplantation but the other died of sepsis without any count recovery.

**Figure 3.**
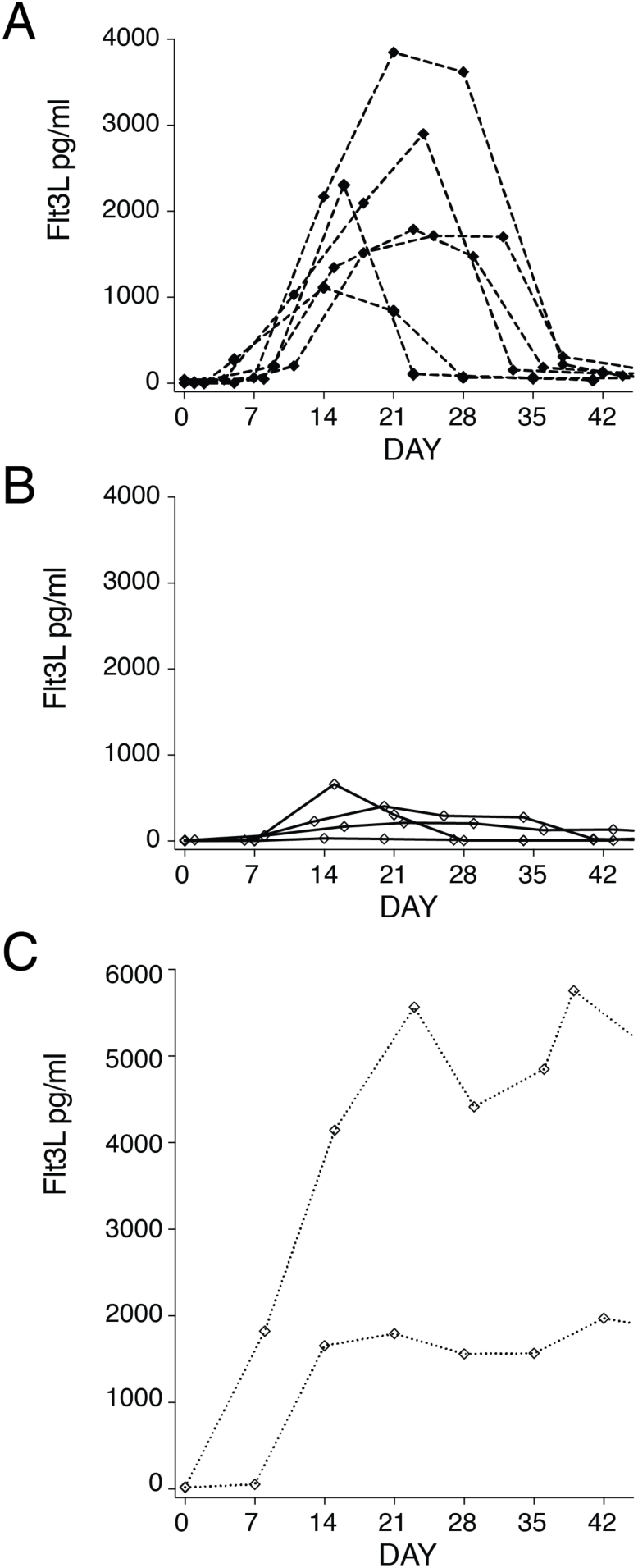
kinetics of Flt3L response during induction. Sequential patients with weekly measurement of Flt3L during induction chemotherapy (n = 12; 4 Cohort A and 8 Cohort B). Dotted line indicates the timing of a single Flt3L measurement in patients treated on the AML17 trial. **A**. Weekly Flt3L in 6 patients entering CR following induction chemotherapy **B**.Weekly Flt3L in 4 patients with refractory disease or partial response following induction chemotherapy **C**.Weekly Flt3L in 2 patients rendered aplastic by induction chemotherapy

### Analysis of AML17 CR and survival according to course 1 Flt3L

Weekly sampling in 12 patients suggested that the response of Flt3L to chemotherapy might predict attainment of CR. In order to test this in a larger cohort, we re-analysed Flt3L levels at day 26 of induction recorded in 140 patients with Flt3 ITD or kinase domain mutation who were treated within the UK NCRI AML17 trial. In these patients, a single Flt3L measurement was performed at day 26 of induction, following 10 days of chemotherapy and 14 days of either tyrosine kinase inhibitor lestaurtinib or placebo ^23^.We first analysed Flt3L in relation to attainment of CR and found a significantly lower median level in patients not achieving CR, including 4 patients with undetectable levels (**Figure 4A**). A receiver-operating characteristic identified an optimal cut-point of 116 pg/ml with AUC of 0.735, sensitivity of 0.721 and specificity of 0.737. There was no difference in response according to lestaurtinib exposure (not shown).

**Figure 4.**
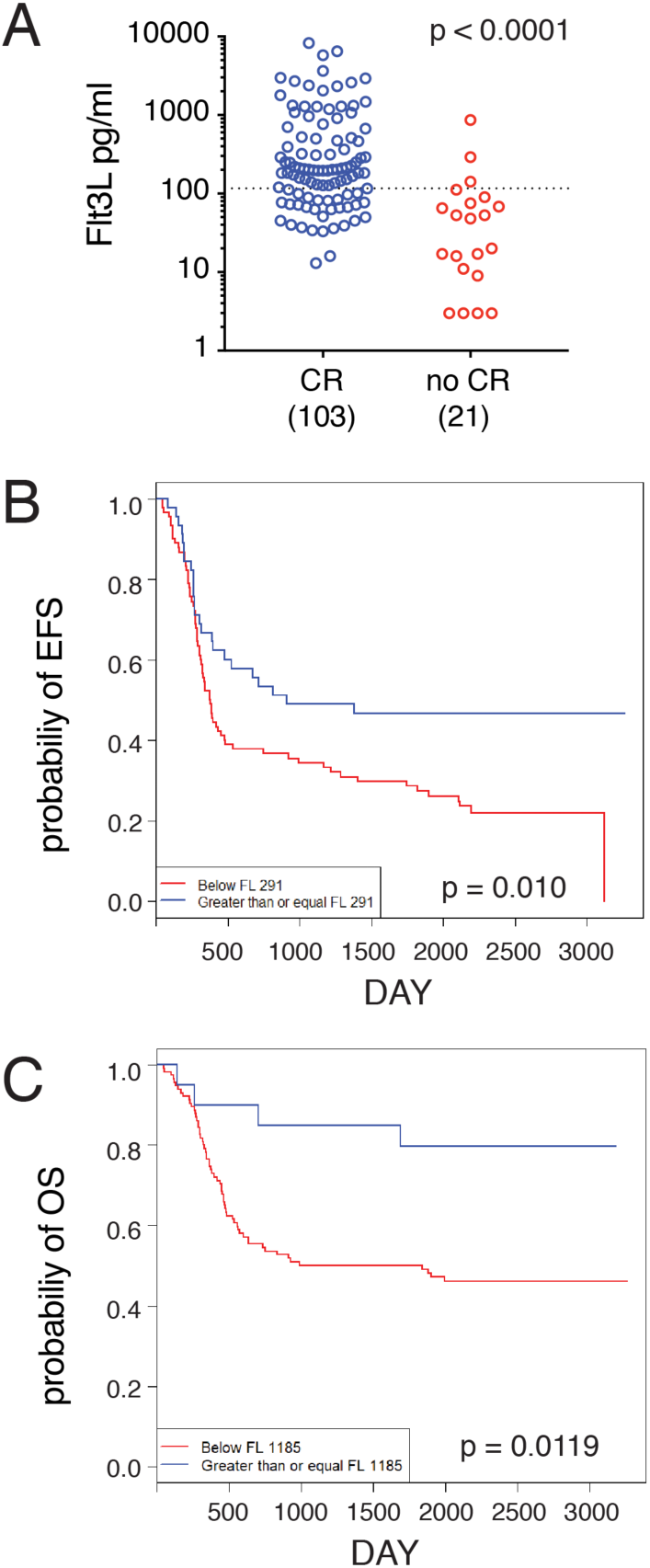
Association between Flt3L, post course 1 remission and survival. Serum Flt3L was measured on day 26 of induction in non APL AML patients with Flt3 mutation randomized to lestaurtinib (day 11-24) on the UK NCRI AML17 trial **A**.Serum Flt3L at day 26 in patients who subsequently achieved or did not achieve CR. Dotted line indicates optimal cutpoint identified by ROC characteristic: 116 pg/ml. Mann Whitney p value **B**.Event-free survival of subgroups with serum Flt3L level <291pg/ml or ≥291pg/ml. Events include: failure to achieve CR; relapse or death from any cause. Threshold of Flt3L for maximum discrimination identified by log rank testing. Log rank p value. **C**.Overall survival of subgroups with serum Flt3L level ≥1185pg/ml or <1185pg. Threshold of Flt3L for maximum discrimination identified by log rank testing. Log rank p value.

This analysis suggested that Flt3L measured at a single point on day 26 had a modest ability to predict CR status post course 1. Consistent with the previous data from Cohort A **(Figure 2)**, none of the patients with undetectable Flt3L at day 26 achieved CR. In practice, these patients may be salvaged by the second course of chemotherapy, as also illustrated in Figure 2, so there is not necessarily an association between Flt3L measured during the first course and more distal outcomes such as EFS and OS. In order to test this, a maximally selected log-rank test was used to define cut points for Flt3L in relation to EFS and OS. This determined a significantly greater EFS for patients with Flt3L above 291pg/ml at day 26 (log rank p = 0.010; **Figure 4B)** and an OS advantage for patients with Flt3L greater than 1185 pg/ml (log-rank p value 0.0119; **Figure 4C)**.

### Post-transplant remission assessment using Flt3L

In order to assess the potential utility of Flt3L in identifying patients at risk of relapse following hematopoietic stem cell transplantation (HSCT) samples were collated from a calendar-driven prospective serum collection from day −7 to 24 months from patients receiving HSCT in Cohort A. From this were identified 8 sequential patients who suffered relapse and 7 patients in continuing CR over the same period (**Figure 5**). In all patients, Flt3L increased to a peak of approximately 1,000pg/ml during the aplastic phase, immediately following conditioning. Flt3L remained at or above physiological levels in 7 patients with sustained remission (**Figure 5A**), but declined and became undetectable in 8 patients who relapsed (**Figure 5B**). Samples were only collected every 3 months so it was not always possible to observe a decline in Flt-3L prior to relapse. However, in 6/17 post-transplant relapses, a drop in Flt3L occurred at least 10 days prior to the diagnosis of relapse.

**Figure 5.**
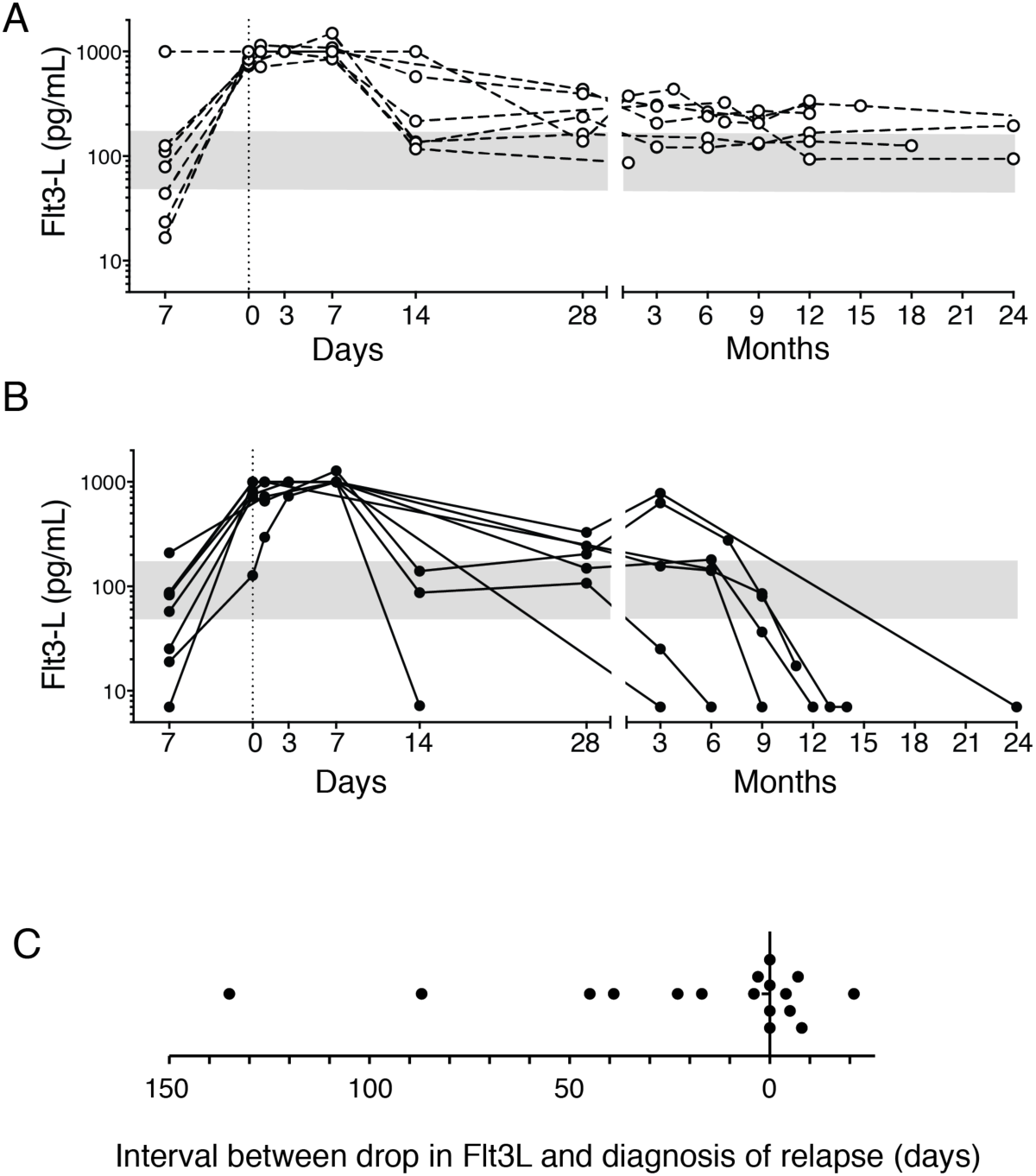
Serial measurement of Flt3L post hematopoietic stem cell transplantation. **A**.Serial serum Flt3L measurements in patients with non-APL AML undergoing hematopoietic stem cell transplantation, remaining in con continuous remission (n = 8; open circles and broken lines). Patients were sampled prior to conditioning (day - 7), on the day of transplant (day 0) and on days and months post-transplant, as indicated. Patients were selected over the same period as B. **B**.Serial serum Flt3L measurements in 7 patients with non-APL AML undergoing hematopoietic stem cell transplantation who relapsed (n = 7; filled circles and broken lines). Consecutive relapsed patients up to 2 years post-transplant, are reported. **C**.Interval between drop in Flt3L below the normal range and diagnosis of relapse. Summary of calendar-driven Flt3L measurement in 17 patients, indicating timing of the point at which Flt3L first went below normal, relative to the hematological diagnosis of relapse. Includes all patients in B with 10 additional patients in whom at least 1 Flt3L measurement was recording post-transplantation.

## Discussion

Flt3L is a critical hematopoietic growth factor that becomes markedly elevated in response to depletion of the hematopoietic stem/progenitor compartment. Here we show that it is depleted in the overwhelming majority of patients with AML, the exceptions being associated with low expression of the receptor Flt3 (CD135) by blasts, as frequently observed in APL. These results are consistent with a model in which the level of Flt3L is inversely related to the abundance of Flt3^+^progenitor cells. Several previous reports have highlighted the expression of Flt3 on AML blasts ^25, 26^and documented that Flt3L may function as an autocrine growth factor ^27^and possibly attenuate the effect of Flt3 inhibitors during therapy ^10^.

Regulation of Flt3L production is not completely understood. In the steady state, the major source appears to be non-hematopoietic ^28^but it is also secreted by activated and peripheral blood T cells ^17, 28^. Alternative splicing produces membrane-bound Flt3L that may play a major role in homeostasis but was not studied here ^29^. Preformed Flt3L has also been described in progenitor cells and AML blasts ^30^. In mice, Flt3L has been described mainly in the hematopoiesis of dendritic cells ^31^but in humans, Flt3 is expressed by CD38^low^CD45RA^+^early progenitor cells suggesting a more fundamental role in the regulation of lympho-myeloid hematopoiesis ^11, 32^.

The potential utility of Flt3L to assist in the assessment of remission between courses of chemotherapy was demonstrated by restoration of Flt3L as patients entered remission, but not in the setting of refractory disease. Morphological assessment and Flt3L were not always concordant, however; 5 patients thought not to be in CR after course 1 showed elevations of Flt3L that heralded attainment of morphological CR after course 2. In 3 cases, CR was reported but there was no restoration of Flt3L and early relapse ensued. These cases suggest that Flt3L might add value to the remission assessment but further prospective study will be required using more accurate standards of remission, such as MRD status by flow cytometry or genomics.

Weekly Flt3L measurement provided a fascinating insight into the dynamics of response to cytoreductive chemotherapy. Patients achieving CR had significantly elevated Flt3L peaking just after the end of therapy and returning to the normal range upon count recovery. Those with refractory disease showed transient elevation of Flt3L followed by rapid depletion. Two patients who were rendered aplastic maintained high levels in the absence of further chemotherapy. Based on these observations, serum ELISA for Flt3L appears to be a low-cost non-invasive means for serial monitoring of patients during remission induction. In particular, Flt3L is likely to dichotomize patients with delayed count recovery into those with low Flt3L due to residual disease and those with high Flt3L and hypoplasia. Tracking of the Flt3L response may thus potentially reduce the need for serial invasive BM assessments and stratify patients for further therapy.

The development of Flt3L as a prognostic tool is more challenging. We took advantage of AML17 trial data in which a single measurement was performed at day 26. A strong association with attainment of CR was observed in the expected direction, at a cut point of 116 pg/ml. This is approximately the median of the normal range, reinforcing the rationale of Flt3L as a biomarker of progenitor cell mass. A more accurate predictive tool might be developed from serial weekly measurements in which the rate of rise, peak level or AUC could have higher discriminatory potential. Concerning longer term survival outcomes, we found discriminatory thresholds for EFS and OS. EFS was inferior in patients with Flt3L below 291pg/ml. Failure to achieve CR was included as an event so this subgroup includes all the patients with refractory disease, who did not survive. A subgroup of 20 patients with high Flt3L (>1185 pg/ml) also enjoyed unusually good overall survival of more than 80% at 3 years, compared with approximately 50% in the remainder of the cohort. The reason for this is not immediately apparent. We speculate that very high levels of Flt3L occur in association with deep remissions in chemo-sensitive disease but further studies will be required to validate these thresholds and determine their prognostic utility in the context of minimal residual disease assessment. We note that all patients were taken from a relatively homogeneous genetic group with either Flt3 ITD or kinase domain mutation. Consequently, we found no evidence to suggest that the elite survivor subgroup group had unusually favorable genetics.

Finally, we explored the use of Flt3L measurement to detect relapse in patients post hematopoietic stem cell transplantation. As expected, there was elevation of Flt3L during aplasia and a recovery to normal or supranormal in patients in continuing CR. Using a 3 monthly sampling schedule it was possible to detect relapse early in a proportion of patients. As with post-chemotherapy evaluation, Flt3L may provide useful insights into pancytopenia in the post-transplant patient, distinguishing between poor graft function or myelosuppression and incipient relapse. The temporal resolution of this would clearly be improved by weekly sampling in future studies. Monitoring for progression or relapse is not limited to transplant patients; two patients with MDS in which Flt3L became undetectable as they progressed to AML, illustrate that Flt3L could facilitate the timing of surveillance bone marrow testing in this patient group.

Remission assessment after the first course of induction chemotherapy is a key milestone in achieving long-term relapse-free survival of AML. Presently, the standard of care is expectant and remission is formally assessed by examination of the bone marrow following hematopoietic recovery. Patients will not know that they have resistant or refractory disease until several weeks after completing the first course of chemotherapy, a problem that is frequently compounded by delayed blood count recovery in those most at risk of treatment failure.

Treatment refractoriness is difficult to predict from clinical, cytogenetic and molecular genetic information ^3, 4^. An interim bone marrow assessment at day 15 may discern blast clearance and is routine practice in many North American centers ^33^. However, this is often a nadir for the patient in terms of cytopenia and attendant systemic illness and interpretation can be difficult. It is not a universal practice in Europe ^7^.

The data presented here suggest that Flt3L may provide an inexpensive, rapid and non-invasive means of assessing remission status in AML. As with MRD techniques, it is likely that performance of the test will be specific to the intensity of treatment. For example, a different kinetic of response, has previously been reported in the context of lower intensity treatment with azacytidine and sorafenib ^34^.

Finally, the response of Flt3L to chemotherapy is likely to reflect multiple known risk factors in AML including genetic subgroup, blast count and attainment of MRD-negative status. Large prospective studies will be required to evaluate this information in a stratified manner in order to determine whether Flt3L responses are integrative of known risk factors or provide independent prognostic information. Both have the potential to find utility in clinical practice.

## Acknowledgements

We thank Hyun Yu (Newcastle University) for assistance with sample collection, Robert Hills (Oxford University) for preliminary statistical analysis and Paresh Vyas (Oxford University) for helpful discussion. C W-B and SK are supported by Cancer Research UK (CRUK) Cardiff Experimental Cancer Medicine Centre core funding and CRUK Clinical Research Committee Clinical Trials Unit Programmme Award core funding. PM and MC are supported by CRUK (C30484/A21025), Histiocytosis UK, the Histiocytosis Association and Bright Red. We also acknowledge support from NIHR Newcastle Biomedical Research Centre at Newcastle upon Tyne Hospitals NHS Foundation Trust.

## Author contributions

Paul Milne: designed the study, acquired, analyzed and interpreted data and drafted the manuscript.

Charlotte Wilhelm-Benartzi analyzed and interpreted data and performed critical revision.

Mike Grunwald acquired and analyzed data

Venetia Bigley acquired, analyzed and interpreted data

Amy Publicover acquired and analyzed data

Sarah Pagan acquired and analyzed data

Helen Marr acquired and analyzed data

Gail Jones acquired and analyzed data

Anne Dickinson acquired data

Angela Grech acquired data

Alan Burnett acquired data

Nigel Russell interpreted data and performed critical revision of the manuscript.

Mark Levis conceived and designed the study and interpreted data

Steven Knapper conceived and designed the study, interpreted data and performed critical revision

Matthew Collin conceived and designed the study, analyzed and interpreted data, drafted the manuscript and performed critical revision

---

**Table S1.**
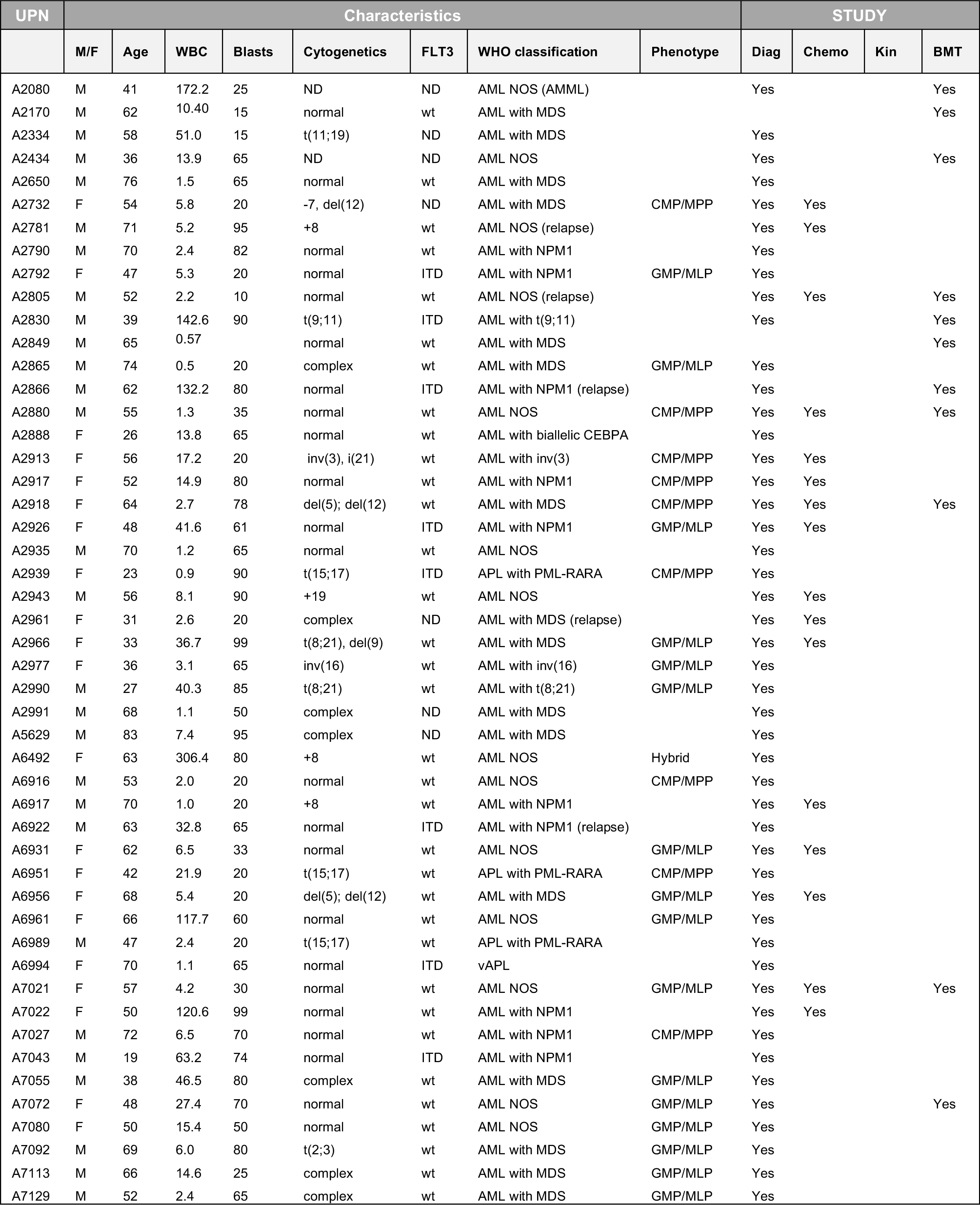
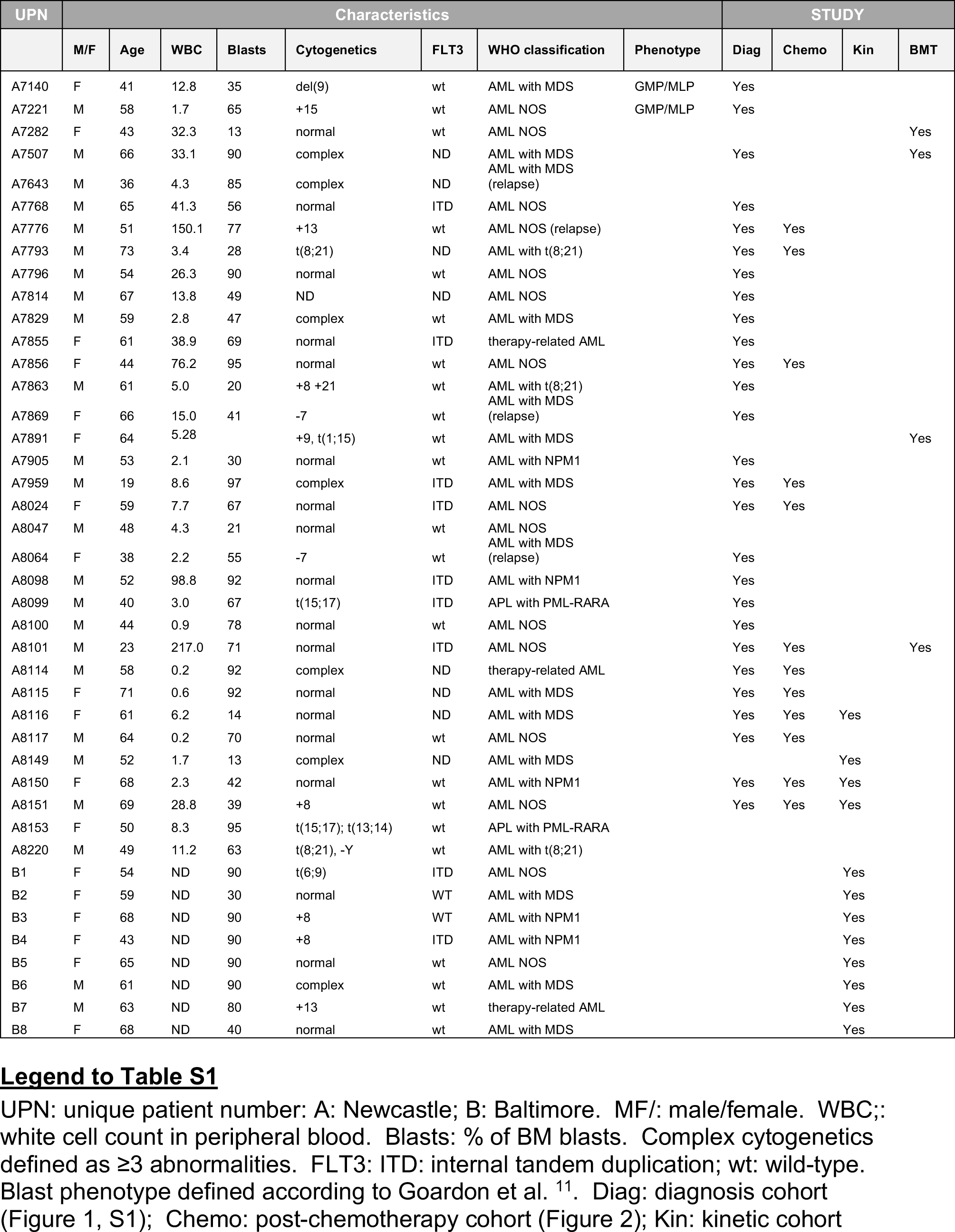
Legend to Table S1 UPN: unique patient number: A: Newcastle; B: Baltimore. MF/: male/female. WBC;: white cell count in peripheral blood. Blasts: % of BM blasts. Complex cytogenetics defined as ≥3 abnormalities. FLT3: ITD: internal tandem duplication; wt: wild-type. Blast phenotype defined according to Goardon et al. ^11^. Diag: diagnosis cohort (Figure 1, S1); Chemo: post-chemotherapy cohort (Figure 2); Kin: kinetic cohort (Figure 3); BMT: bone marrow transplant cohort (Figure 5). Blank cells indicate test or intervention not done.

**Figure S1.**
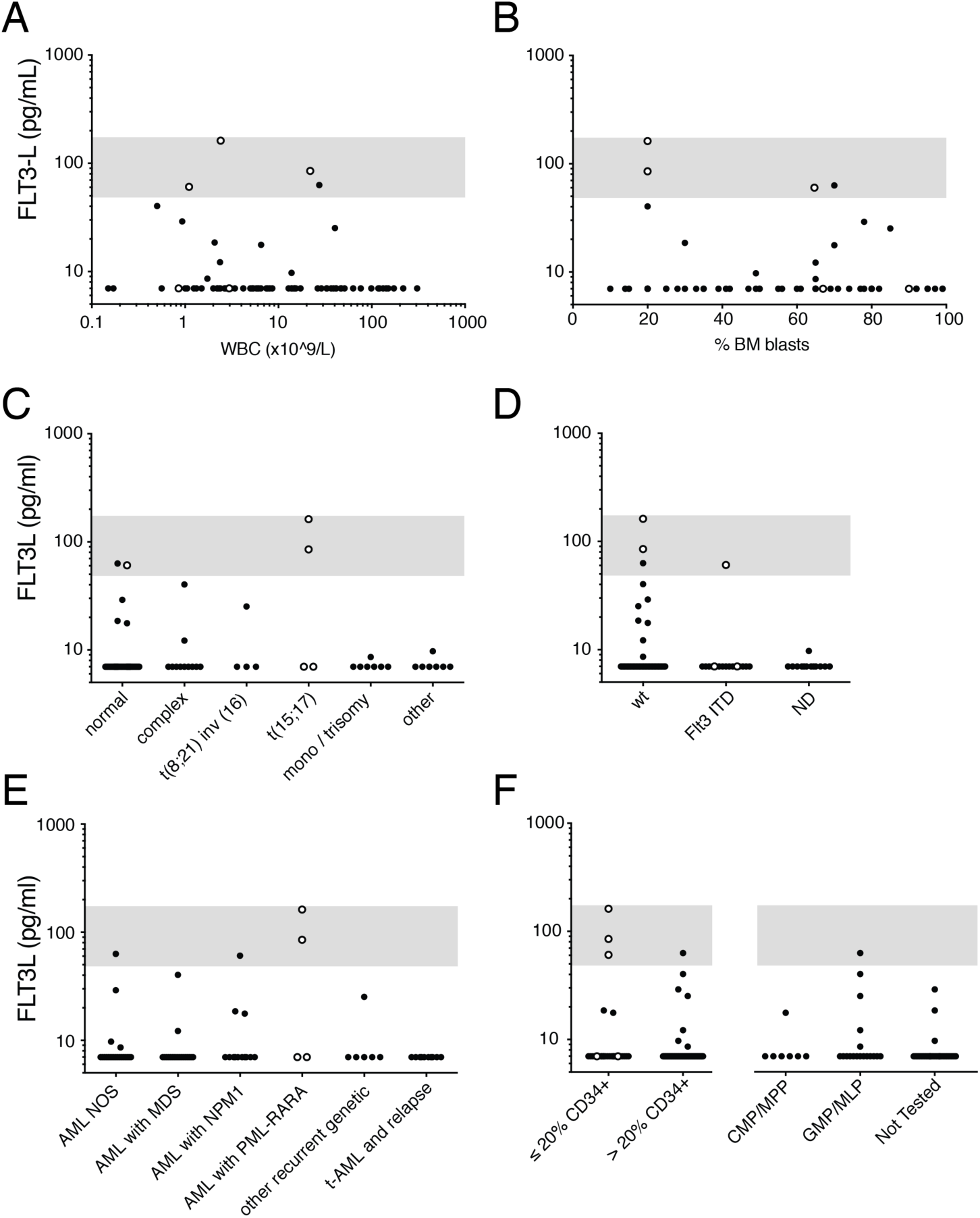
Relationships between Flt3L and other variables at diagnosis. APL cases are shown in open circles (including 1 case of variant APL with typical phenotype and morphology but no detectable PML-RARA rearrangement) **A**.WBC at diagnosis measured by automated counter **B**.% BM blasts estimated from aspirate smear and microscopy **C**.Cytogenetic category **D**.Presence of Flt3 ITD **E**.WHO category **F**.Expression of CD34 and phenotype of CD34+ AML according to Goardon et al. ^11^

